# A Two-Process Model for Circadian and Sleep-dependent Modulation of Pain Sensitivity

**DOI:** 10.1101/098319

**Authors:** Natalia Toporikova, Megan Hastings Hagenauer, Paige Ferguson, Victoria Booth

## Abstract

Pain sensitivity is strongly modulated by time of day and by prior sleep behavior. These two factors, governed by the circadian rhythm and homeostatic sleep drive, respectively, likewise dictate the timing and duration of sleep. The fields of sleep and circadian research have identified much of the physiology underlying the circadian rhythm and homeostatic sleep drive with mathematical modeling playing an important role in understanding how these two processes interact to affect sleep behavior. We hypothesize that the daily rhythm of pain sensitivity and its sleep-dependent modulation reflect an interaction of the circadian rhythm and homeostatic sleep drive. To investigate this hypothesis, we adapt the formalism of a classic mathematical model for the regulation of sleep behavior by the circadian rhythm and homeostatic sleep drive, called the Two Process model, to simulate the interaction of these two processes on pain sensitivity. To construct the model, we utilize data from experimental reports on the daily rhythmicity of pain sensitivity in humans to define a “daily pain sensitivity” function. We decompose this function into two processes: a sleep-dependent process *S*(*t*) that follows the homeostatic sleep drive and a circadian process *C*(*t*) that is dictated by the circadian rhythm. By simulating different sleep schedules with the original Two Process model, we compute changes in the sleep-dependent process *S*(*t*) that modulates pain sensitivity. By combining *S*(*t*) with the circadian process *C*(*t*), our model predicts resultant changes in the daily pain sensitivity rhythm. We illustrate model predictions for changes in pain sensitivity due to sleep deprivation, sleep restriction and shift work schedules. We believe that this model may be a useful tool for pain management by providing predictions of the variations in pain sensitivity due to changing sleep schedules.

## 1 Introduction

Pain sensitivity is strongly modulated by time of day and by prior sleep behavior. As reviewed in the preceding chapter [7], in humans highest sensitivity to painful stimuli occurs during the night and lowest sensitivity occurs in the late afternoon. Additionally, sleep deprivation increases pain sensitivity. Experimental studies of pain sensitivity have only considered either time-of-day (circadian) effects or sleep-dependent effects. However, it is well known that the effects of prior sleep behavior and the circadian rhythmicity of sleep propensity interact to govern the timing and duration of sleep. For example, sleep deprivation causes an increase in the drive for sleep and can promote the occurrence of sleep during daytime hours. However, the circadian rhythm acts to promote wakefulness during the day. The interaction of these two processes, namely the homeostatic sleep drive and the circadian rhythm, results in limited durations of daytime sleep episodes despite elevated sleep drive levels which would prolong sleep if it occurred during the evening.

We hypothesize that the 24 hour and sleep-dependent modulation of pain sensitivity may likewise reflect a combined interaction of these two processes: circadian rhythm and homeostatic sleep drive. As an example, consider a study on the effects of sleep deprivation on pain sensitivity. If pain measurements are conducted in the late afternoon, effects of sleep deprivation may be underestimated, while they may be overestimated if measurements are taken during the night because of circadian modulation of pain sensitivity. To investigate this hypothesis, we adapt the formalism of a classic and influential mathematical model for the regulation of sleep behavior by the circadian rhythm and homeostatic sleep drive, called the Two Process model [3], to model how the interaction of these two processes may affect pain sensitivity.

## 2 Background: Two Process Model for circadian modulation of sleep timing

The original Two Process model for sleep regulation was constructed to account for the interaction of the homeostatic sleep drive and the circadian rhythm in the timing and duration of sleep [3]. While the physiology of the homeostatic sleep drive has not been completely determined [9, 6, 12], a biological marker for it has been identified as the power of low frequency, delta range (0.5 – 4 Hz) oscillations in EEG recordings during human sleep in the non-rapid eye movement (nREM) or slow wave stage of sleep. Specifically, at the initiation of a sleep episode, the power of delta oscillations is high and decays roughly exponentially as sleep continues through the night. These observations motivated modeling the homeostatic sleep drive, Process S 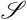 (*t*) in the model, as an exponential function that decreases during sleep and increases during wake. The time constant for the decay during sleep was fit to the decay in delta oscillation power observed in sleep EEG recordings. The time constant for the increase of the homeostatic sleep drive during wake was constrained to match the fit to data during sleep. Specifically, Process S increases exponentially during wake with time constant *τ_w_* = 18.2 h and decreases exponentially during sleep with time constant *τ_s_* = 4.2 h. In wake, 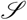(*t*) is governed by:

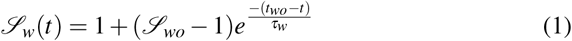

where time *t* is in hours, 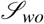 is set to the 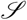 value at the previous wake onset (*wo*) and *t_wo_* is the time of the previous wake onset. In sleep,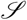(*t*) is governed by:

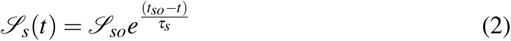

where 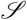*_so_* is set to the 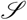 value at the previous sleep onset (*so*) and *t_so_* is the time of the previous sleep onset.

While more is known about the physiology of the circadian rhythm that modulates sleep timing [10, 5], the Two Process model focuses on time-of-day effects on sleep propensity to determine the equations for Process C 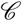(*t*). Experiments measuring typical sleep latencies and durations across the day suggested that sleep timing follows a skewed sinusoidal function such that sleep is minimal near midday and is strongly promoted in the early morning hours [3]. This circadian rhythmicity in sleep propensity is combined with Process S by Process C dictating the threshold values at which Process S transitions from sleep to wake and vice versa. As such, Process C consists of two sinusoidally varying functions 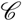_*w*_(*t*) and 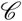_*s*_(*t*) such that 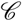*_w_*(*t*) dictates the 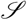(*t*) values when the transition from wake to sleep should occur and 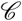*_s_* (*t*) dictates the 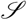 (*t*) values at which sleep to wake transitions occur:

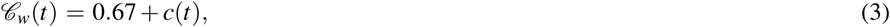

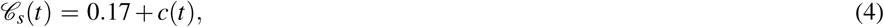

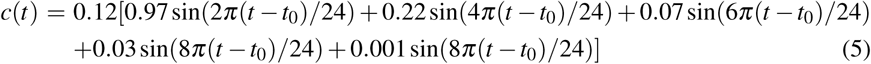

where *t*_0_ sets the circadian phase at the initial time.

With these parameter values, the model generates a 24 h cycle of sleep-wake behavior with 16 h in wake and 8 h in sleep (Figure 1). We initialize *t*, *c* and 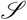 so that the model starts at *t* = 6 for 6am at the beginning of a wake episode with the following initial conditions: *t* = 6, *t*_0_ = 7.5, 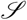 = 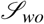 = 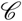*_s_*(6) and *t_wo_* = 6. With these initial conditions, the model simulates a sleep schedule with wake onset at 6am and sleep onset at 10pm. The wake state occurs during the interval that 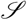(*t*) is increasing and the sleep state occurs when it is decreasing.

**Fig. 1.**
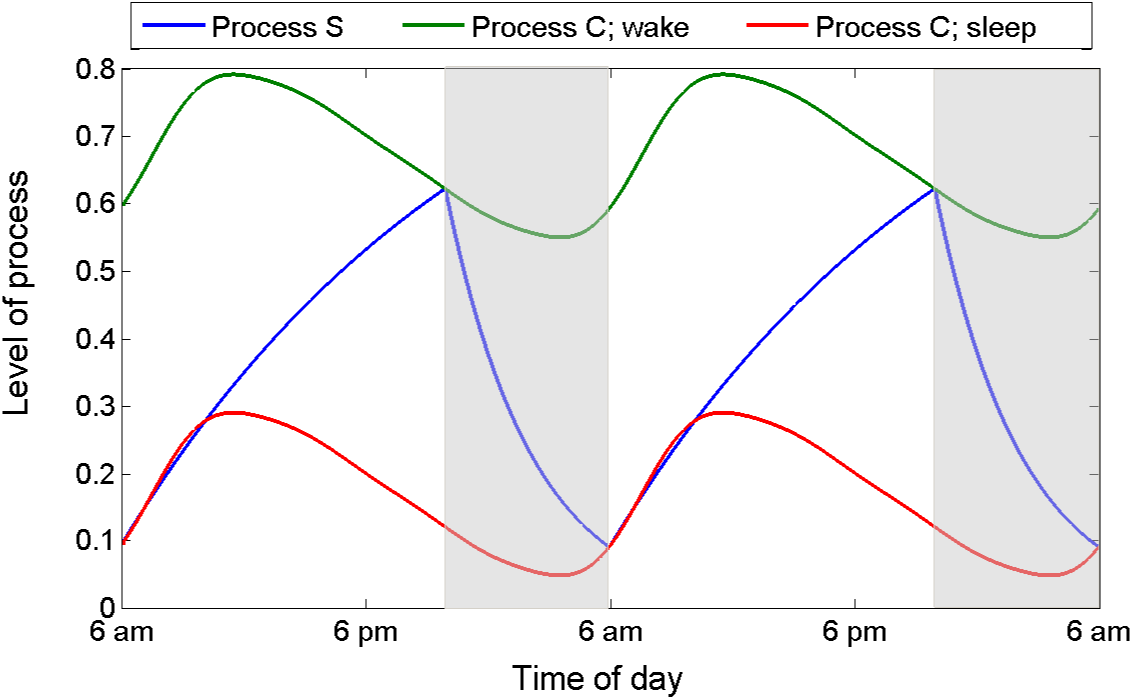
Two Process model predicting the timing and duration of sleep and wake episodes as governed by the homeostatic sleep drive (Process S, 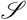(*t*), blue curve) whose levels for transitions between wake and sleep are dictated by the circadian rhythm (Process C, green and red curves). Wake occurs as Process S is increasing with sleep initiated when Process S intersects 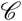*_w_*(*t*) (green curve). Sleep (shaded regions) continues as Process S decreases until it intersects 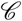*_s_*(*t*) (red curve)

## 3 Two Process Model for Pain Sensitivity

While there is strong experimental evidence that increased homeostatic sleep drive that occurs due to sleep deprivation increases pain sensitivity (as reviewed in [7]), the correlation between pain sensitivity and the homeostatic sleep drive under normal sleep conditions has not been determined. To construct our model, we assume that pain sensitivity exhibits increases and decreases throughout the 24 h day that follow the increases and decreases of the homeostatic sleep drive. In particular, we assume pain sensitivity increases with time spent awake even under normal daily schedules of sleep-wake behavior. We implement the simplest form of this assumption by assuming that sleep dependent modulation of pain sensitivity follows the exponential increase during wake and decrease during sleep of the homeostatic sleep drive predicted by the original Two Process Model. A consequence of this primary assumption is the further assumption that the experimentally observed daily rhythm of pain sensitivity is the combined result of circadian and sleep-dependent modulation. For our model, we develop an estimate of the sleep-dependent component of this modulation by determining an appropriate scaling factor of Process S from the original Two Process Model. We then estimate the circadian modulation of pain sensitivity by subtracting the estimated sleep-dependent component from the experimentally observed daily rhythm of pain sensitivity. We assume that this circadian modulatory component is not affected by sleep behavior. To implement the model, we use the Two Process model to compute variations of the sleep-dependent modulatory component under different patterns of sleep behavior and combine it with the circadian component to predict their combined effect on pain sensitivity with changes in sleep behavior.

As discussed in the companion article [7], multiple studies report a consistent daily rhythm of pain sensitivity that peaks during the night and is at a minimum during the late afternoon. In order to quantify this rhythm, in [7] we constructed a prototypical “daily pain sensitivity function” by normalizing data from four studies that tested experimentally induced pain responses across 24 h. For each dataset, we transformed time to relation to morning wake time and transformed units to percent of the mean of the reported daily variation. Following these transformations, the data collectively formed a tight, sinusoidal curve with a trough ~ 9 h after usual wake onset and a peak ~ 2 h after usual sleep onset (see Figure 1 in [7]). This constructed curve provides the qualitative shape of the daily rhythm in pain sensitivity, but does not accurately reflect the amplitude of the rhythm due to the units transformation. In order to compare the amplitude of effects across different studies on the daily fluctuation of pain sensitivity using different pain modalities, and additionally to compare effects due to sleep deprivation, in [7] we identified new normalizations for units of change in pain sensitivity. Namely, for the particular pain modality used, we converted the observed changes in pain threshold to a percentage of the range of physiologically-meaningful stimulation values or to a percentage of the range of painful stimulation values. Using these units for changes in pain sensitivity, we found that the average amplitude (max - min) of the daily rhythm measured in multiple pain modalities was 12 – 14% (see Table 6 in [7]). Thus, for our model, we define the experimentally observed daily pain rhythm, *P_obs_*(*t*), as the “daily pain sensitivity function” scaled so that its oscillation amplitude (peak - trough) matches this average amplitude. The appropriate scaling parameters for the “daily pain sensitivity function” were in the range [0.6,0.7]. In the model, this curve is assumed to be the combined result of circadian and sleep-dependent modulation of pain sensitivity and will be decomposed into sleep-dependent and circadian components, *S*(*t*) and *C*(*t*), respectively.

To explicitly define the model, let *S*(*t*) represent the time-varying, sleep-dependent modulation of pain sensitivity that follows the homeostatic sleep drive as predicted by the Two Process model:

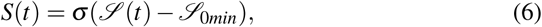

where 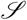(*t*) is Process S (Eqs 1 and 2), 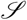_0*min*_ is the minimum value of 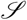 under normal sleep-wake behavior and σ is an appropriate scaling factor. In this way, during normal sleep-wake behavior predicted by the Two Process Model, *S*(*t*) varies between 0 at times of wake onset to a maximum value at times of sleep onset. During other sleep patterns, such as sleep deprivation or restriction when the sleep homeostatic drive may be elevated, increases in *S*(*t*) are thus measured relative to its minimum possible value. We define *C*(*t*)as the circadian modulation of pain sensitivity. Then the observed daily rhythm of pain sensitivity *P_obs_*(*t*) is defined as

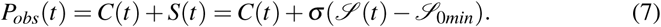

As the physiological processes regulating circadian rhythms are believed to be independent of the processes regulating sleep homeostasis, we assume that *C*(*t*) and *S*(*t*) are likewise regulated independently. Thus, under conditions of sleep deprivation or restriction that do not change the circadian rhythm, we assume that the circadian modulation of pain sensitivity *C*(*t*) does not vary, while the sleep-dependent component *S*(*t*) would vary with variation in Process S. For example, consider the scenario of 8 h sleep deprivation due to extending wake 8 h beyond the normal sleep onset time of 10pm in the Two Process Model. Let *S_SD_*_8_(*t*) = σ(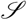_*SD*8_(*t*) – 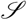_0*min*_) be the modified sleep-dependent pain modulation where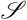 *_SD8_(t)* is Process S under this instance of 8 h sleep deprivation and 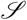_0*min*_ is the minimum value of Process S under normal sleep-wake behavior. As in the original Two Process Model, this sleep disruption is assumed not to significantly affect circadian rhythms, thus the predicted pain sensitivity in this scenario, *P*_*SD*8_(*t*), is given by

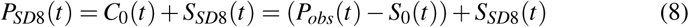

where *S*_0_(*t*) = σ(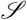_0_(*t*) – 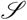_0*min*_) and 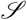_0_(*t*) is Process S under normal sleep-wake behavior.

To identify an appropriate value for the scaling parameter σ, we constrain the model to replicate the change in pain sensitivity observed after one night of sleep deprivation. Specifically, σ is chosen so that *S*(*t*) reaches values between 12 – 14 after 8 h of sleep deprivation as modeled with the Two Process Model to reflect the experimentally observed approximately 13% increase in evoked pain responses measured relative to the range of painful stimulation (see Table 6 in [7]). This yields values for σ in the interval [18.1433,21.1672].

We now compute *C*_0_(*t*) as given in Eq (8) as

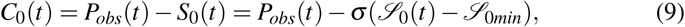

where 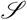_0*min*_ = 0.0953, the minimum value reached by 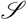_0_(*t*) under normal sleep behavior.

In summary, our model for the predicted rhythm of pain sensitivity due to circadian and sleep-dependent modulation under a specific sleep-wake behavior pattern *α*, *P_α_* (*t*), is defined as

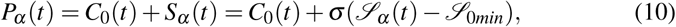

where 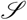*_α_* (*t*) is Process S as simulated by the Two Process model under sleep-wake behavior pattern *α* and and 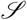_0*min*_ is its minimum under normal sleep-wake behavior. We note that the range of scalings for *P_obs_*(*t*) (namely, [0.6,0.7]) and σ ([18.1433,21.1672]), reflecting the experimentally reported average effects of daily rhythm and sleep deprivation on evoked pain responses, leads to a range of predicted values for *C*_0_(*t*), *S_α_*(*t*) and *P_α_*(*t*) which are indicated by the thickness of the curves in the middle and bottom panels of Figures 2 – 4.

**Fig. 2.**
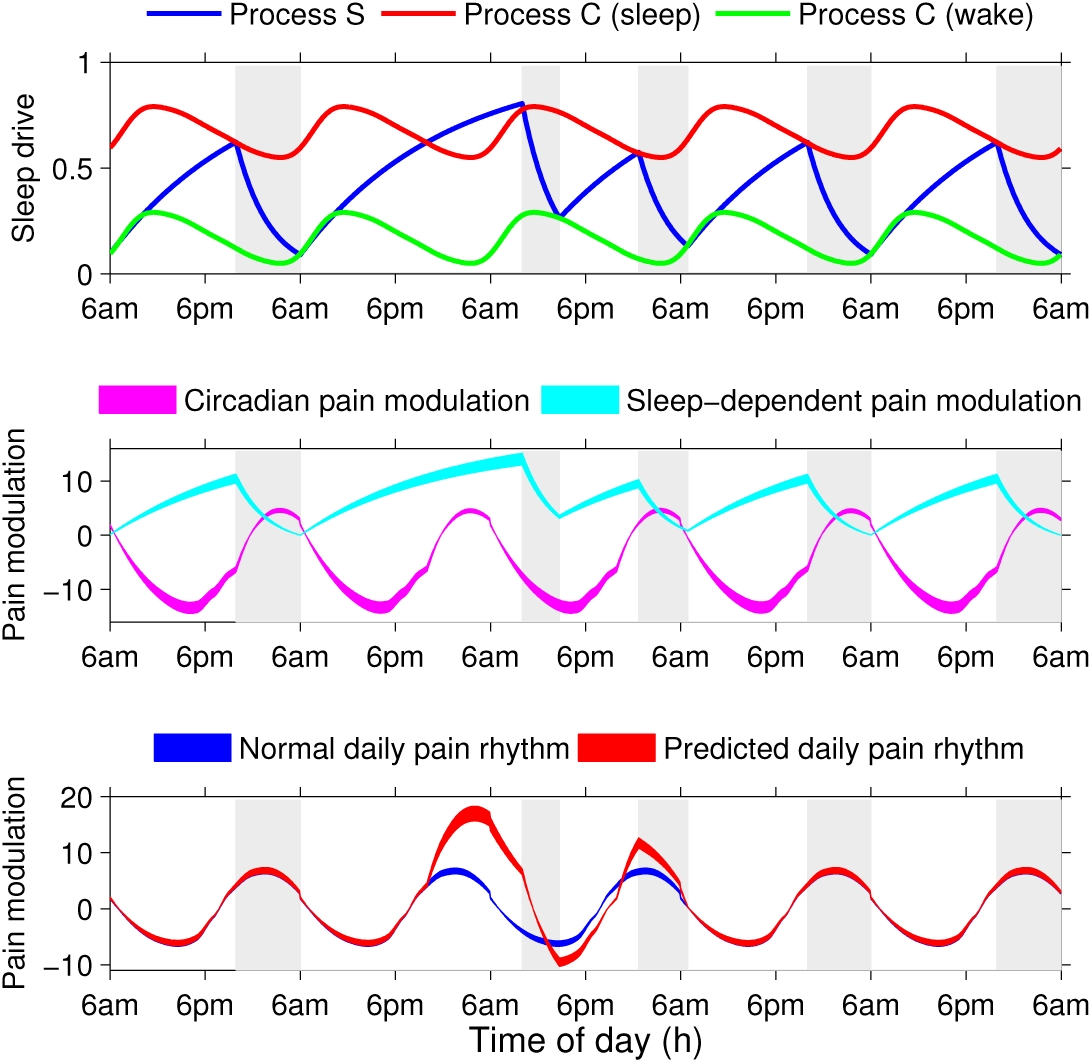
Predicted pain sensitivity during a simulation of continuous sleep deprivation protocol for 28 hours. Shaded regions represent sleep times. Top: Simulation of the Two Process model with 12 hours of sleep deprivation initiated at the customary sleep onset time on day 2. Middle: The sleep-dependent pain modulation (cyan curve) increased during the sleep deprivation protocol while the circadian pain modulation was unperturbed (magenta curve). Bottom: Combining the sleep-dependent and circadian pain modulation yielded the predicted daily rhythm in pain sensitivity (red curve) that showed increases in sensitivity during the deprivation protocol and decreases the following day, compared to the rhythm under the normal sleep schedule (blue curve). The thickness of the curves in the middle and bottom panels indicate the range of values obtained due to the range of scalings for *P_obs_*(*t*) and σ (see Section 3). Vertical axis units for top: level of homeostatic sleep drive normalized to be between 0 and 1; for middle and bottom: levels of sleep-dependent and circadian pain modulation are set so that the normal daily pain rhythm (blue curve) has amplitude 12-14 with mean 0.

**Fig. 3.**
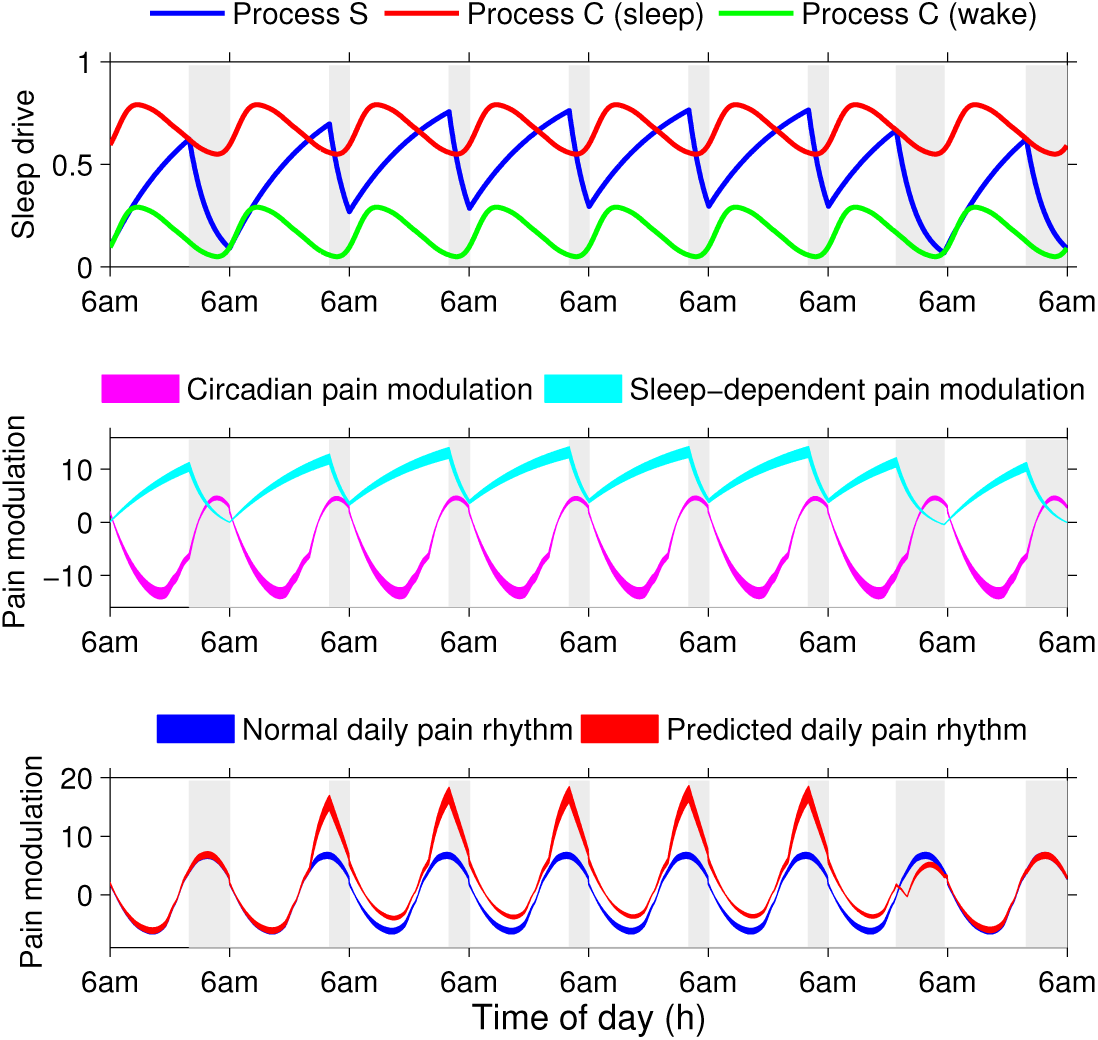
Predicted pain sensitivity during a simulation of sleep restriction to 4 hours a night for 5 consecutive nights. Shaded regions represent sleep times. Top: Simulation of the Two Process model with sleep restricted to occur between 2 — 6 am for days 2 — 6. Middle: Sleep-dependent pain modulation (cyan curve) remained elevated for days 2 — 6 while the circadian pain modulation was unperturbed (magenta curve). Bottom: The predicted daily pain sensitivity (red curve) showed nonuniform increases across the days when sleep was restricted, compared to levels under the normal sleep schedule (blue curve). Vertical axis units are the same as in Figure 2.

**Fig. 4.**
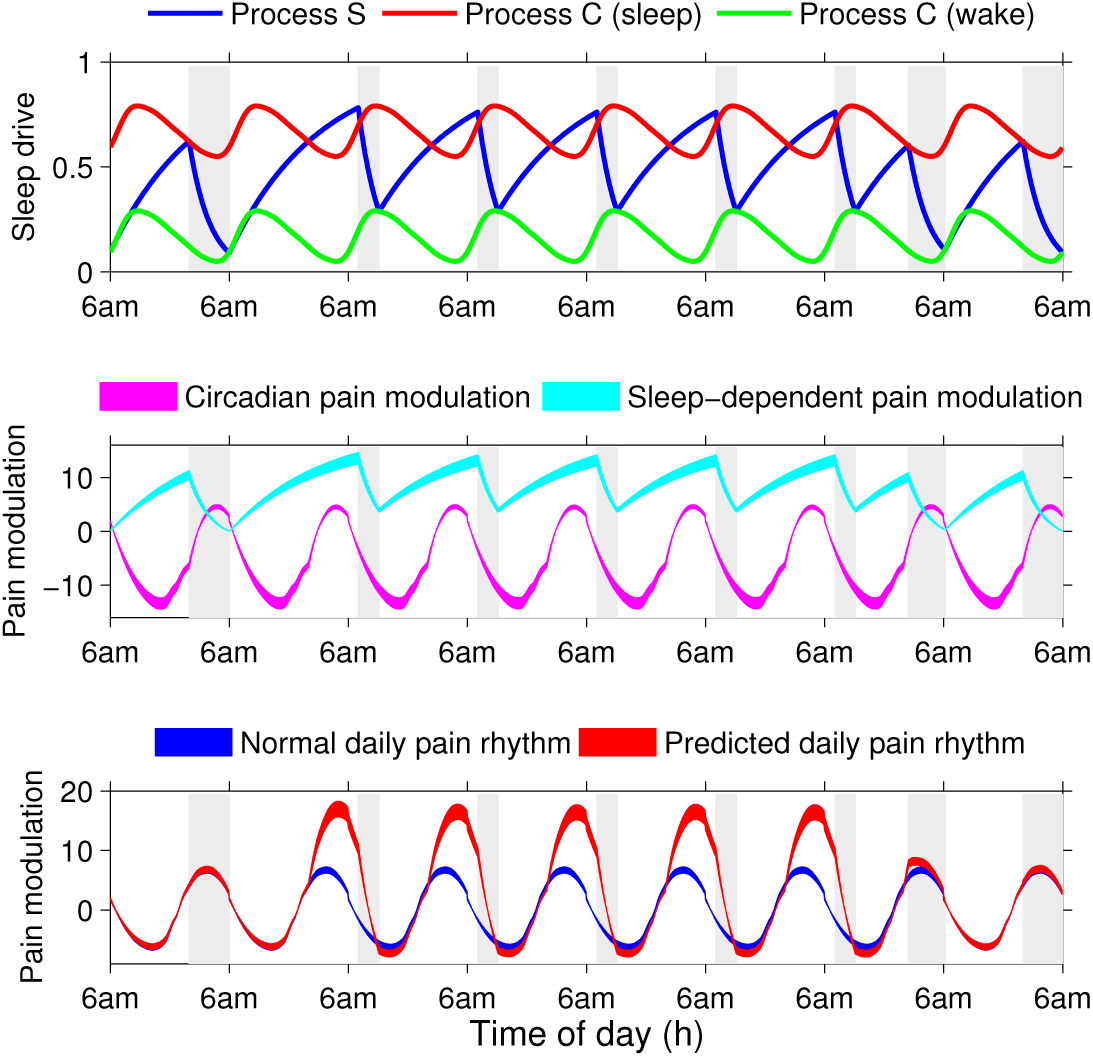
Predicted pain sensitivity under simulated shift work schedule from 11 pm to 7 am for 5 consecutive days. Shaded regions represent sleep times. Top: Simulation of the Two Process model with sleep onset suspended until 8 am on days 3-7. Middle: Sleep-dependent pain modulation (cyan curve) remained elevated for days 2-6 while the circadian pain modulation was unperturbed (magenta curve). Bottom: The predicted pain sensitivity (red curve) showed significant increases during the majority of the shift work hours and during sleep episodes, compared to levels under the normal sleep schedule (blue curve). Vertical axis units are the same as in Figure 2.

## 4 Model predictions

To predict how sensitivity to pain was affected by sleep schedules, we applied our model to three different simulations of disrupted sleep, namely sleep deprivation, sleep restriction and a shift work schedule. To simulate the effect of these sleep schedule perturbations on the homeostatic sleep drive, we manually induced sleep or wake transitions in the Two Process model and ignored the state transition thresholds dictated by Process C. At the end of the sleep perturbation protocol, we re-initiated the Process C threshold crossing rules for sleep initiation and termination.

### 4.1 Pain sensitivity under sleep deprivation

First, we tested the change in pain sensitivity due to a continuous sleep deprivation protocol (Figure 2). This numerical experiment simulated a common protocol in which human subjects are kept awake for 12 h beyond their customary bed time. To obtain the behavior of Process S in this protocol, 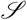*_α_*(*t*) = 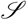_*SD*12_(*t*) in Equation 10, we ran the Two Process model for 5 days during which the first 24 h simulated the customary wake time at 6 am and sleep initiation at 10 pm. On day 2 of the simulation, we initiated the continuous sleep deprivation protocol, by ignoring the threshold crossing condition for Process C for 28 hours. After 28 hours, we reinitiated the rule for sleep initiation when Process S is above 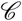*_W_*, which resulted in immediate sleep onset. For the remainder of the simulation, the Two Process model followed its usual evolution (Figure 2, top panel).

During the sleep deprivation protocol, Process S continued to increase exponentially beyond its usual values (top panel, blue curve), which drove an increase in sleep-dependent modulation of pain, *S_α_* (*t*) = *S*_*SD*12_(*t*) in Equation 10 (middle panel, cyan curve). When *S*_*SD*12_(*t*) was added to the circadian modulation of pain function,*C*_0_(*t*) (middle panel, magenta curve), the resulting predicted pain sensitivity function, *P_α_*(*t*) = *P*_*SD*12_(*t*) in Equation 10 (lower panel, red curve) showed a large increase in sensitivity, compared to normal levels (lower panel, blue curve), during the deprivation period which continued into the subsequent sleep episode. An interesting prediction of the model is a reduction in pain sensitivity due to the sleep deprivation. Right after awakening from the recovery sleep episode, pain sensitivity was lower than under the normal sleep schedule. This occurs because of a phase shift in the perturbed sleep-dependent pain modulation, whose minimum coincides with the minimum of the circadian pain modulation. This reduction reversed as the circadian modulation neared its peak because the model remains in waking beyond the normal sleep onset time, leading to an increase in sleep-dependent pain modulation.

### 4.2 Pain sensitivity under sleep restriction

Another common sleep disruption that occurs in people's daily lives and is induced in experimental settings is sleep restriction during which the time allowed for sleep is restricted over several consecutive days. We simulated a sleep restriction protocol in which sleep is allowed for only 4 hours per night, during 2 — 6am, for 5 consecutive nights as might occur in a typical work week (Figure 3). We simulated the Two Process model for 8 days where on days 2 — 6 sleep onset and wake onset were manually induced for the sleep restriction protocol (top panel) to obtain 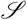*_α_*(*t*) = 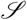_*SR*4_(*t*) (top panel). Following Process S, the sleep-dependent pain modulation *S_α_* (*t*) = *S*_*SR*4_ (*t*) was elevated during days 2 — 6 (middle panel, cyan curve) predicting continuous increased pain sensitivity over those days (bottom panel, red curve). The interaction of the sleep-dependent and circadian pain modulation led to non-uniform increases in pain sensitivity across the day. The smallest increase occurred during the evening hours after 6 pm, but pain sensitivity increased significantly during the hours between the customary sleep onset time of 10 pm and the allowed sleep onset time of 2 am. During the allowed sleep episode, pain sensitivity decreased but remained elevated at wake onset at an intermediate level. On day 7 when sleep was allowed to occur normally, pain sensitivity decreased during the sleep episode as the homeostatic sleep drive decayed to normal levels.

### 4.3 Pain sensitivity under shift work schedules

Shift work and the resulting misalignment of sleep schedules with the circadian rhythm have been associated with a myriad of adverse health conditions, including increased rates of obesity, cardiac disease and cancer [8, 18]. Additionally, shift work has been correlated with increased rates of reported musculoskeletal and lower back pain [17, 15, 1]. We simulated an 8-h shift work schedule between 11 pm and 7 am for 5 consecutive days as may occur for a typical shiftwork week (Figure 4). Sleep onset was assumed at 8am to allow for commuting time between work and home. We simulated the Two Process model (top panel) for 8 days during which on day 2 sleep onset was suspended until 8 am on the morning of day 3. The duration of sleep was allowed to be dictated by the model. Subsequent shifts were simulated by again suspending sleep onset until 8 am on the morning of the following day. After each shift, sleep behavior was allowed to resume as predicted by the model. This simulation of the Two Process model generated 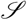*_α_* (*t*) = 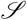*_SW_* (*t*) from which the sleep-dependent pain modulation *S_α_* (*t*) = *S_SW_* (*t*) was computed (middle panel, cyan curve). The predicted pain sensitivity (bottom panel, red curve) was significantly elevated during the majority of time during the work shift and during the subsequent sleep episode. A decrease in pain sensitivity occurred for a short duration upon awakening due to a decrease in sleep-dependent pain modulation when the circadian pain modulation was near its minimum.

## 5 Conclusions and Future Work

We have developed a mathematical model to investigate the combined influences and interactions of the homeostatic sleep drive and the circadian rhythm on the variation of pain sensitivity across the day. While experimental results indicate that these two processes have strong effects on pain sensitivity, their combined influences, especially under conditions of disrupted or limited sleep behavior, have not been explored. The model adapts the formalism of a successful mathematical model for the regulation of sleep behavior by the circadian rhythm and homeostatic sleep drive, called the Two Process model [3]. Similar to the Two Process model, our model for pain sensitivity *P*(*t*) assumes that the sensitivity level is independently modulated by a circadian process *C*_0_(*t*) and by a sleep-dependent process *S*(*t*). The effects of these two processes were derived from the experimentally observed daily rhythm of pain, *P_obs_*(*t*), computed from multiple experimental results, by applying the assumption that the sleep-dependent process *S*(*t*) is driven by the homeostatic sleep drive as predicted by the Two Process model. The resulting circadian modulation *C*_0_(*t*) imparts a roughly sinusoidal modulation to *P*(*t*) that peaks during the early morning hours before customary awakening and has a trough in the late afternoon hours (middle panels of Figures 2 – 4, magenta curves). The sleep-dependent modulation *S*(*t*) exponentially increases during wake and decreases during sleep (cyan curves).

The strength of the model comes from the ability to predict how the daily cycle of pain sensitivity will change due to changes in sleep behavior. The original Two Process model has been validated against many diverse disrupted sleep schedules, such as sleep deprivation and sleep restriction. Our model incorporates these results of the Two Process model to predict the consequent variations in pain sensitivity, as illustrated here for 12 h of sleep deprivation, 4 h sleep restriction and shift work schedules.

Another advantage of the model is the ability to quantitatively compare the effects of different sleep schedules on pain sensitivity. For example, the simulated shift work schedule shown in Figure 4 results in 4.1 h of sleep on the 5 shift work days, approximately 30 minutes more total sleep time as during the simulated restricted sleep schedule shown in Figure 3. Despite similar amounts of sleep during these two schedules, there are large differences in the effect of these schedules on pain sensitivity. The shift work schedule resulted in higher maximum pain sensitivity, inducing an increase equal to 98.7% of the maximum amplitude of the daily observed rhythm in pain sensitivity *P_obs_*(*t*), while the sleep restriction schedule induced a 78.9% increase. The timing of this maximum also differed with it occurring 2 h prior to sleep onset in the shift work schedule and at sleep onset in the sleep restriction schedule. Thus, while the shift work schedule allows for slightly more sleep, it results in higher peak pain sensitivity that may be more apparent or debilitating since it occurs during active waking periods compared to the sleep restriction schedule. However, the sleep restriction schedule results in a larger portion of the waking period with increased pain sensitivity compared to the shift work schedule. Under shift work, during approximately 70% of the waking period there are negligible changes in pain sensitivity while under sleep restriction pain sensitivity is increased throughout the entire waking period. This type of quantitative comparison possible with the model would be especially useful in evaluating potential sleep interventions, such as nap schedules, in alleviating pain sensitivity increases.

Our model assumes that pain sensitivity is determined by the homeostatic sleep drive. This assumption is based on findings from sleep deprivation experiments, however additional experiments are needed to validate it. Sleep drive is believed to be due to some structure that tracks or some substance that accumulates a 'need to sleep' during prolonged wakefulness, and discharges this homeostatic need during sleep. Several neuromodulators/neurotransmitters have been proposed to serve a role of a homeostatic accumulator for the need to sleep, with adenosine as a leading candidate[2, 14]. During prolonged wakefulness, adenosine levels rise in some sleep-related areas of the brain [11]. Injection of adenosine causes sleep in cats and rats [13,14]. Hence, at least one mechanism for homeostatic sleep drive might be an accumulation of adenosine that enhances the activity of sleep-promoting brain areas and reduces activity in wake-promoting brain areas. Therefore, a test of our modeling assumption that the homeostatic sleep drive increases pain sensitivity would be to investigate changes in pain sensitivity in animals injected with adenosine. If our assumption is correct, pain sensitivity will increase after the adenosine injection.

Patients with chronic pain often suffer from insomnia and report pain as the primary reason for their disrupted sleep. Our model assumes that the circadian rhythm and homeostatic sleep drive determine daily cycles of pain sensitivity. However, other physiological or cognitive factors surely affect responses to painful stimuli. For example, the perception of painful stimuli might be affected by its timing relative to wake onset. While cognitive functions are reduced for a short time after waking from sleep, a phenomenon known as sleep inertia [16], this cognitive state has interesting effects on the perception of pain. It has been shown that sleep inertia has no effect on pain perception when subjects are awoken abruptly from slow wave or non-rapid eye movement (non-REM) sleep but it reduces pain perception when awoken abruptly from REM sleep [4]. While the reasons for this difference is not known, these results suggest that patients who wake up in pain either perceive accurately the pain they are experiencing, or at worst underestimate the level of pain if woken from REM sleep.

The model is phenomenological in nature; specific physiological mechanisms underlying sleep-dependent and circadian modulation of pain are not explicitly modeled. Instead, the pain sensitivity variable of the model has units of percent change in sensitivity relative to the estimated range of painful stimulation for the particular pain modality, and we've shifted it so that the mean sensitivity is zero. This phenomenological form has the advantage that the model can be validated against data for different modalities and measures of pain sensitivity. In this way, we believe the model has the potential to be a useful tool for pain management by providing predictions of the variations in pain sensitivity due to changing sleep schedules. Future work will focus on identifying data sets for model validation. The advent of activity monitoring devices, such as fitbit (fitbit.com), that can continuously track sleep behavior, and wearables that can prompt users to easily record pain sensitivity at multiple time points per day would provide the type of continuous data on wake and sleep schedules and pain that would be ideal for further developing our model.

## Acknowledgements

The authors thank Anita Layton, the organizer of A Research Collaboration Workshop for Women in Mathematical Biology during which this work was initiated and the other participants of the workshop for valuable feedback and discussion, especially Jennifer Crodelle and Sofia Piltz. This work was conducted as a part of A Research Collaboration Workshop for Women in Mathematical Biology at the National Institute for Mathematical and Biological Synthesis, sponsored by the National Science Foundation through NSF Award DBI-1300426, with additional support from The University of Tennessee, Knoxville. VB acknowledges the support of NSF Award DMS-1412119.

